# Insight into biogeochemical models from Scale Transition Theory: A dimensionless, scale-free approach

**DOI:** 10.1101/2020.04.13.039818

**Authors:** Chris H. Wilson, Stefan Gerber

## Abstract

Leading an effective response to the accelerating crisis of anthropogenic climate change will require improved understanding of global carbon cycling. A critical source of uncertainty in Earth Systems Models (ESMs) is the role of microbes in mediating both the formation and decomposition of soil organic matter, and hence in determining patterns of CO2 efflux. Traditionally, ESMs model carbon turnover as a first order process impacted primarily by abiotic factors, whereas contemporary biogeochemical models often explicitly represent the microbial biomass and enzyme pools as the active agents of decomposition. However, the combination of non-linear microbial kinetics and ecological heterogeneity across space guarantees that upscaled dyamics will violate mean-field assumptions via Jensen’s Inequality. Violations of mean-field assumptions mean that parameter estimates from models fit to upscaled data (e.g. eddy covariance towers) are likely systematically biased. Here we present a generic mathematical analysis of upscaled michaelis-menten kinetics, grounded in Scale Transition Theory. We advance the framework by providing solutions in dimensionless form, and illustrate how this approach facilitates qualitative insight into the significance of this scale transition, and argue that it will facilitate future cross site intercomparisons of scale transition effects from flux data. We also discuss the critical terms that need to be constrained in order to unbias parameter estimates.

## Introduction

The current crisis of anthropogenic climate change is expected to accelerate during the 21st century. Despite considerable effort to better constrain global biogeochemical models, considerable uncertainty remains about how best to represent emerging mechanistic understanding of soil element cycling into process-based models (Wieder et al. 2015, 2018; Todd-Brown, Zheng, and Crowther 2018). This is a critical gap in knowledge because variations among models predict hugely varying responses to global change drivers such as temperature, soil moisture, and *CO*_2_ enrichment. For example, a traditional first-order linear model forecasts no change or even slight enhancement of SOC pools by 2100 whereas one microbial-explicit model forecasts a loss of ∼70Pg of C, depending on whether microbial physiology acclimates to higher temperatures (Wieder, Bonan, and Allison 2013). In general, our understanding of how carbon (and other elements) cycles in soil is undergoing significant revision toward a more microbial-centric paradigm. In contrast to traditional first-order linear models (e.g. CENTURY, Parton et al. (1987)), microbial explicit models feature non-linear dynamics in which microbial biomass (or, similarly, microbially-driven enzyme pools) are responsible for decomposition, in addition to providing substrate for synthesis of potentially long-term SOC (Blankinship and Schimel 2018; Blankinship et al. 2018). While indisputably a better representation of our scientific knowledge, non-linear microbial models face several well-known challenges, including less analytical tractability, greater computational challenges, and uncertainty about structural formulation and dynamics (Georgiou et al. 2017; Sihi et al. 2016). However, one critical consequence of non-linear microbial models that is only recently gaining attention is their implications for addressing the upscaling challenge.

While the fields of population and community ecology have long confronted the challenges posed by non-linearity and heterogeneity in spatiotemporal scaling of ecological dynamics (Chesson 2009; Levin 1992), ecosystem ecology and biogeochemistry have tended to approach the challenge of scale either by 1) utilizing mean-field assumptions, or 2) addressing the challenge of scaling via grid-based computational/numeric methods. While there is nothing wrong inherently with either approach, they unfortunately cannot yield theoretical insight into the consequences of non-linearity and heterogeneity for scaling. Briefly, the combination of non-linearity and heterogeneity means that aggregated behavior differs systematically from mean-field predictions, a special case of Jensen’s Inequality. In mathematical notation:

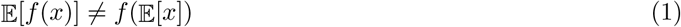

Although Jensen’s Inequality is well-known from basic probability theory (Ross 2002), it’s implications for ecological dynamics under heterogeneity were not well-appreciated until the pioneering work of Peter Chesson in the 1990s (Chesson 1998). In the case of carbon cycle science, there are a few immediate and critical applications. For instance, most trace gas emission processes are well-known to be non-linear functions of underlying drivers such as temperature and soil moisture. For example, ecosystem respiration (hereafter “*R*_*eco*_”) is an exponential function of temperature (usually expressed in *Q*_10_), and a unimodal function of soil moisture. Thus, when matching observations of *CO*_2_ efflux (“*F* “) to ecosystems, variations in soil temperature and moisture could imply that *F* differs systematically from a mean-field prediction. In addition to missing critical analytical insight, not accounting for this behavior might have severe consequences for inverse modeling and estimation of the parameters governing process-based models (PBMs). The basic consequences of Jensen’s Inequality for estimation of trace gas emission (*CH*_4_ and *N*_2_*O*) were first discussed by Van Oijen et al. (2017), but have not been picked up on elsewhere, until the present work (and very recently by Chakrawal et al. (2019)).

Chakrawal et al. (2019) provide a detailed and compelling first-pass application of scale transition theory to biogeochemical modeling. Our contribution here complements their laudable effort by providing a more generic mathematical analysis of the scale transition, equally applicable to both forward and reverse michaelis-menten microbial kinetics. As in Chakrawal et al. (2019), we address the consequences of heterogeneity in both substrate/microbes (“biochemical heterogeneity”) as well as in the kinetic parameters (“ecological heterogeneity”). However, we diverge from their approach in that, rather than explore detailed simulation models, we derive a completely non-dimensionalized expression for aggregating non-linear microbial kinetics over both types of heterogeneity simultaneously. We illustrate the enormous clarity this brings in several special cases of our full analysis. Altogether, our approach provides universal insight into the properties of the scale transition, and enables clear conclusions to be drawn across systems in terms of the role of spatial variances and covariances in shaping ecosystem carbon efflux. Our work provides a simplified, yet systematic framework around which to base subsequent empirical and simulation-based studies.

### Carbon Efflux and the Scale Transition

#### Setup

A variety of microbial-explicit PBMs have been proposed in the literature, starting with the classic enzyme pool model of Schimel and Weintraub (2003). In order to elucidate universal properties of the scale transition, we focus here on the *CO*_2_ efflux following decomposition of a single substrate by a single microbial pool obeying Michaelis-Menten (MM) dynamics:

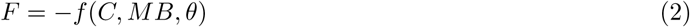

where *F* is the *CO*_2_ flux, *θ* is a vector of parameters, specifically *V*_*max*_ (the maximum reaction rate given saturation of either C, in forward MM, or MB, in reverse MM), *k*_*h*_ (the half-saturation constant), and *CUE* = 1 − *ϵ* (accounting for carbon-use efficiency).

Our specific model for *F* is:

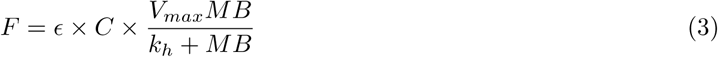

Following the terminology of, the above is our “patch” model and our goal is to understand how spatial variances and covariances impact the integrated flux, which represents the spatial expectation or 𝔼 [*F*] (hereafter denoted 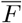), which represents 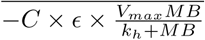.

The incorrect approach to solving for 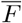 is to simply plug-in the mean-field solution:

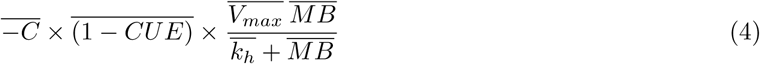

Analytically, an exact solution would require specification of a joint distribution for C and MB - *π*(*MB, C, θ*) - and solution of the convolution integral:

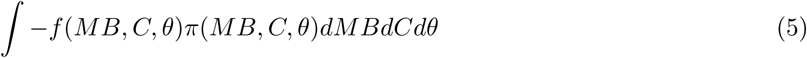

However, following Chesson (2012) and Chakrawal et al. (2019) we are free to approximate the solution for arbitrary distributions using a Taylor Series approximation expanded to the 2nd moment. Specifically, we take the expectation over a multivariable Taylor Series expansion, centered around the mean-field values of all parameters *θ* (for simplicitiy, the variables *MB* and *C* are included in the parameter vector *θ*):

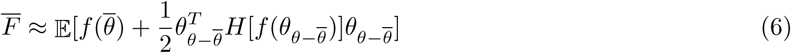

where *H*[*f* (*θ*)] represents the Hessian matrix of the function that determines the *CO*_2_ efflux *F* (in this case Michaelis-Menten), 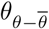 represents the deviation from the mean at each instance and for each of the parameters. It can easily be seen that 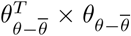 is the variance-covariance matrix, and that the first moment of the Taylor expansion cancels because the first derivative of 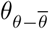 is zero.

### Non-dimensionalization

Expanding equation 6 out, we have 5 terms involving the variances of *C, MB*, 1 − *CUE, V*_*max*_, and *k*_*h*_, and 10 terms involving covariances among the parameters. We can redistribute the expectation operator over this approximation to see that we are dealing with the contributions from the variance-covariance terms, weighted by the second partial derivatives evaluated at the mean for each parameter. However the resulting expression **does not readily yield insight into the impact of scale transition upon the dynamics**, since second partial derivatives and cross partial derivatives do not have easy intuition. Morever, variances and covariances depend arbitrarily upon the scale of units and measurements involved, hindering both intuition and cross-site comparisons. Therefore, we non-dimensionalize equation 6 for 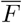 as follows:

1. We define a dimensionless quanity *λ* as 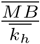. *λ* thus represents a multiplicative factor expressing the ratio of the mean microbial biomass over it’s mean half-saturation value.
2. We divide all of the terms in 6 by their mean-field value, and represent the whole equation as a product:

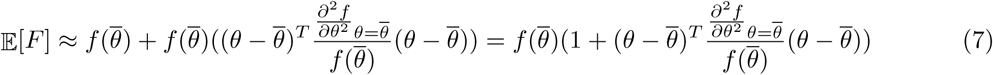
3. We calculate the resulting expression for 𝔼 [*F*]
4. We notice that 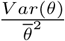 can be re-expressed as 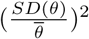 which in turn is the square of the dimensionless coefficient of variation (*CV* (*θ*))^2^. This enables us to reformulate the variance terms in (7).
5. Similarly, since the covariance terms can be rewritten as *COV* (*X, Y*) = *ρ*_*X,Y*_ *SD*(*X*)*SD*(*Y*), we have the following equality:

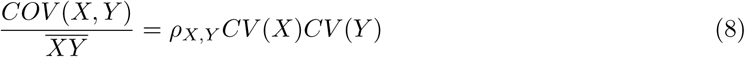

Applying steps 1-5 to all the terms in the equation, we end up with a fully dimensionless equation:

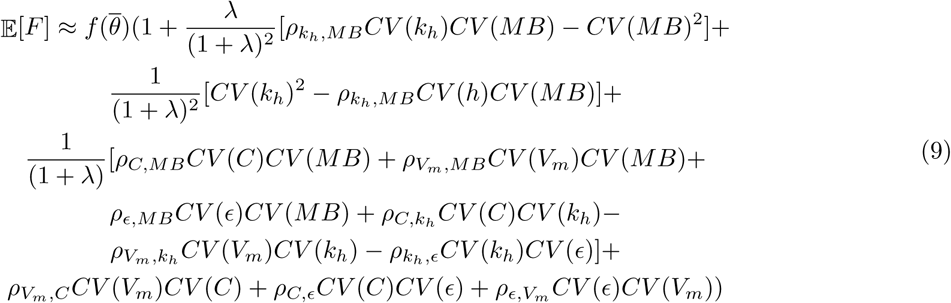

Note that by symmetry, we have also solved for the case of the forward Michaelis-Menten kinetics. This can be expressed simply by interchanging *C* and *MB*, and by correspondingly altering *λ* to represent the ratio of substrate availability over half-saturation.

## Discussion

### Generic Insights

Having fully non-dimensionalized equation 6 into equation 9, we are in a much better position to gain analytical insight into the scale transition. But first, we note that Equation 9 reveals a multitude of terms that must accounted for in understanding how an upscaled flux measurement (F) can violate mean-field assumptions when derived from sites incorporating ecological heterogeneity. Therefore, when ecosystem models are fit to flux data, the relevant parameter estimates are likely to be biased, perhaps severely so. Consider the situation where the sum of variability and correlation terms to the right of the 1 on the RHS are positive, leading to upscaled fluxes systematically larger than mean-field. In this situation, with no correction the tendency will be to inflate the estimates of parameters giving larger fluxes (e.g. *V*_*m*_, *ϵ*). The opposite will occur when the relevant variability and correlation terms are negative.

Analytically, our framework emphasizes the pivotal role played by the quantity *λ* throughout this equation. *λ* scales the contributions of the parameter variation and correlation terms to the deviation from mean field behavior according to the ratios 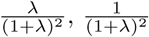 and 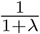. In particular, all of the parameter variance terms (which have become *CV* (*θ*)^2^ upon non-dimensionalization), are scaled by one of these three *λ* ratios, alongside 7 out of 10 of the covariance terms. Overall, low *λ* (here *λ <*≈ 1) keeps all of the spatial correction terms in play, while increasing *λ* tends to simplify matters. As noted by Sihi et al. (2018), as *MB* → ∞ (equivalent to MB » *k*_*h*_ or *λ* → ∞), reverse Michaelis-Menten kinetics converge to first order, leaving:

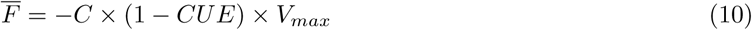

Accordingly, in our setup, the multiplicative factor for the scale transition correction approaches a simplified expression, as *λ* → ∞:

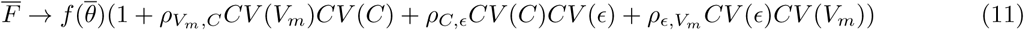

This is actually quite remarkable. Despite invoking the situation where microbial biomass (and its enzyme supply) is effectively infinite - thus linearizing the underlying patch models - we cannot eliminate the possibility of a potentially substantial deviation from mean-field when scaling decomposition kinetics. We note that in this resulting expression, we have reduced the situation to a set of three critical correlations involving two microbial physiological parameters (*ϵ*, and *V*_*m*_), and substrate availability (*C*). Regardless of their respective variabilities (CV terms), if these correlations are close to zero, then the whole expression converges to mean field.

Returning to the situation where *λ* is not large, if we ignore the correlation terms (i.e set them to zero), we see that there are direct contributions to the scale transition from the variability in *MB* and *k*_*h*_ that may, to some extent, balance each other:

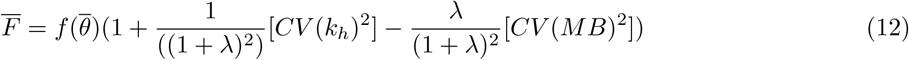

Focusing on the offsetting correction terms, we can re-write as:

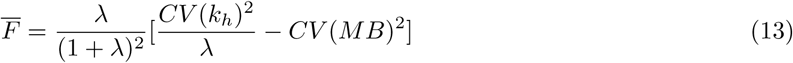

and for the case of *λ* = 1, this becomes:

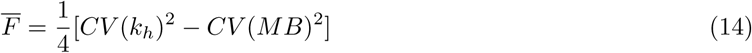

Thus, variability in the factors of soil protection that impact upon *k*_*h*_ in practice, can offset the impact of variability in microbial biomass itself.

More generally, using our dimensionless equation 9 puts modelers and empiricists in a better position to assess the quantitative significance of the scale transition correction across systems compared to expressions with uninterpretable second partial derivatives and cross derivatives, and arbitrarily scaled variance terms. By re-expressing 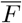 in terms of dimensionless coefficients of variation, correlation coefficients and *λ*, we can plug-in realistic values for variability in any relevant parameter and assess the % effect on 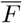 in terms of deviation from mean field behavior. We argue that this formulation possesses significant advantages not only in understanding how to scale flux estimates 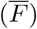 *within* a site, but also is more likely to facilitate intercomparison *among* sites in terms of their scale-free variability.

### Spatial Colocation of Microbes and Substrate

To illustrate these advantages in interpretability, we first take the special case of a model where we treat all parameters as constant (and known) except substrate and microbial biomass. This corresponds to setting the other CV() and *ρ* terms to 0. In this case, we are isolating the impact of the spatial colocation of substrate and decomposers. Our equation for 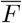 becomes:

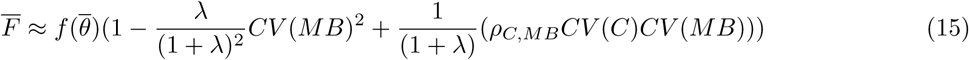

First, we note that the magnitude of the mean field correction scales as the *square* of the coefficient of variation of microbial biomass. Quadratic scaling means that at low to moderate levels of variability, the deviation from mean field behavior is likely to be fairly minimal, but at moderately high to high levels of variability, severe deviations can be expected. For instance, a five-fold increase in *CV* (*MB*) from 0.1 to 0.5 would result in a 25-fold increase in the scale transition factor.

Additionally, we note that, in the case of this formulation, there is a very clear dual convergence as *lambda* increases:

1. deviation from mean-field behavior declines, and
2. first order kinetics are approached

Indeed, our equation 15 reveals the exact speed of this convergence in terms of dimensionless *λ* and a balance of *CV* (*MB*), *CV* (*C*) and their correlation.

In figure 1, we illustrate the scale transition solutions to equation 15 as a function of lambda for various choices of CV(C), CV(MB) and *ρ*:

**Figure 1:**
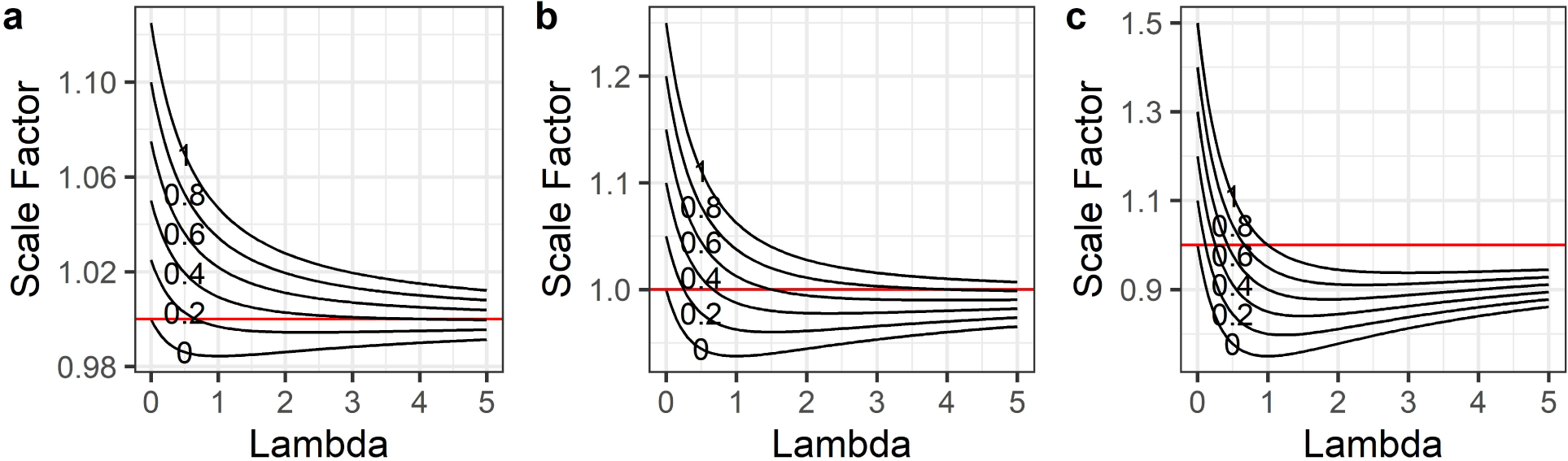
Scale correction factor for models given spatial colocation between microbes and substrate across a gradient of λ values, and for a variety of correlation ρ values (0-1), with CV(SOC) held constant at 0.5, a) CV(MB) = 0.25, b) CV(MB) = 0.5, and c) CV(MB) = 1.

We see in Fig. 1 that in the case of pure spatial colocation, with no variation in the kinetic parameters, the scale transition correction factor varies from a maximum of 1.5 to a minimum around 0.75, and in all cases indeed converges to 1 as lambda increases. The variability assumed for C and MB impacts primarily the scale of the correction factor, and secondarily the qualitative behavior as *λ* and *ρ* vary (Fig. 1a-c).

We argue that the principal virtue of having a simplified, generic dimensionless equation of this sort is that it enables us to think in a unit-free/scale-free manner about the *plausible* range of the scale transition correction given transparent assumptions about the relevant variabilities and correlations. For instance, it should not be a very difficult task in any given study system to estimate the variation in *MB* and *C* and their correlation over space. In principle, we could therefore understand the spatial scale transition arising from colocation over rather large ecosystems, provided these quantities are well-behaved and well-constrained from data. If successful, this achieves two goals: 1. obviates the need to parameterize grid-based numerical models to understand scaled-up fluxes, and 2. facilitates ready comparison among sites given estimates of the dimensionless quantities involved.

In passing, we note that if we introduce a new term *λ*_2_ representing the relationship between *CV* (*MB*) and *CV* (*C*) as follows *CV* (*C*) = *λ*_2_*CV* (*MB*), we can re-express the mean-field correction factor as:

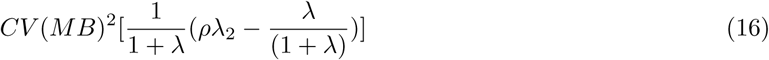

Thus, whether the correction is positive or negative depends crucially on the product of the colocation correlation coefficient *ρ* and the extent of variability in substrate relative to variability in microbes. Setting the inner term in 16 to zero and solving for *ρ*, we find:

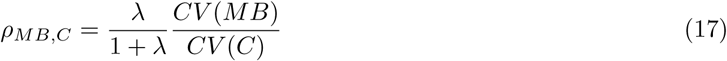

If we fix *λ*_2_ to unity, as done in our Fig.1, and focus interest on solutions around *λ*_1_ = 1 our mean-field correction factor simplifies to:

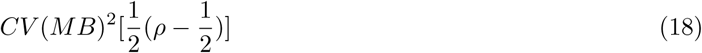

In general, it is clear that the scale transition correction is larger to the extent that substrate variability exceeds microbial variability under reverse michaelis-menten kinetics (the opposite relation holds for forward Michaelis-Menten by symmetry). Thus, *our analysis highlights in equation 17 another route of convergence back to the mean field beyond the simple increase of λ.*

### Environmental Heterogeneity

So far we have analyzed in depth the role of variability in microbes and their substrate, but not in the ecological drivers underlying maximal reaction rates (i.e. *V*_*max*_) or half-saturation (i.e. *k*_*h*_). We start with the observation that both linear first order and non-linear microbial models will show characteristic scale transitions given heterogeneity in temperature and soil moisture. Consider the asymptotic convergence of the patch model for reverse MM to first order as *MB* → ∞:

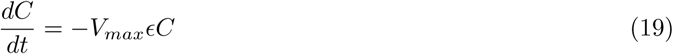

This is mathematically equivalent to the more standard way of writing these models down as:

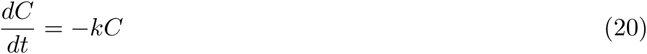

To make matters clear, we re-express the rate limiting maximal reaction velocity *V*_*max*_ first as a function of temperature (assuming all else constant):

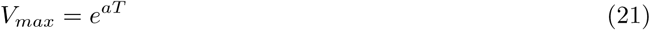

In this case, our integrated flux equation will be:

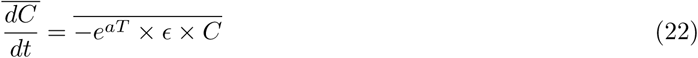

Allowing for variability in *T*, this integrated equation will show characteristic scale transitions given the convex (exponential) relationship with *T*.

Using the Taylor expansion again to second order we have:

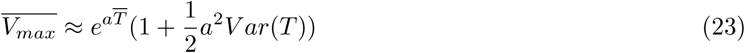

The critical scale correction term here is again multiplicative, and we re-express it into a function of a dimensionless coefficient of variation parameter more suited to ready interpretation. First, the exponential dependence of respiration on temperature is canonically codified in terms of *Q*_10_ scaling. We substitute 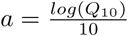, and end up with:

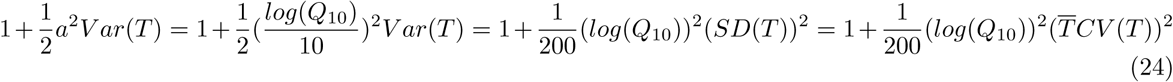

For a “typical” *Q*_10_ of 2.5, and a 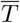 of 25, we see the multiplicative scale transition correction in figure 2:

**Figure 2:**
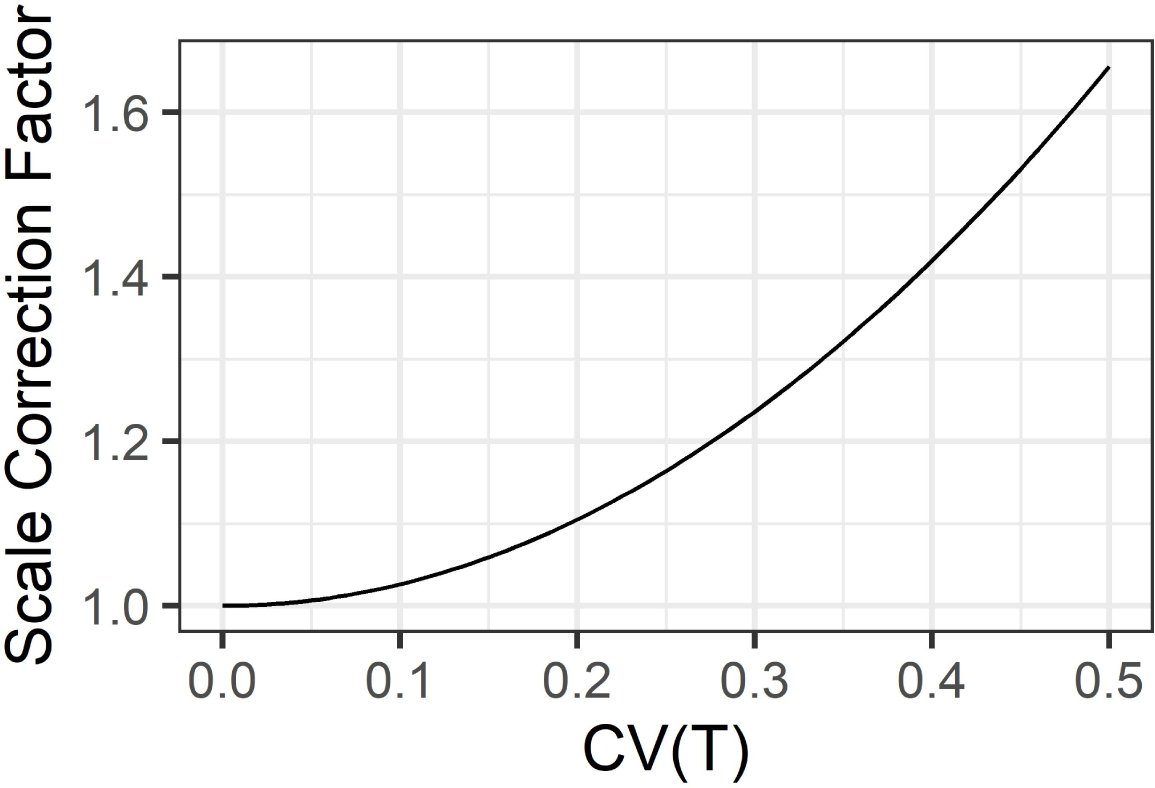
Scale correction factor for Q10 temperature response scaling given coefficient of variation CV(Temp) from 0 to 0.5, 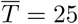, and Q10=2.5

**Figure 3:**
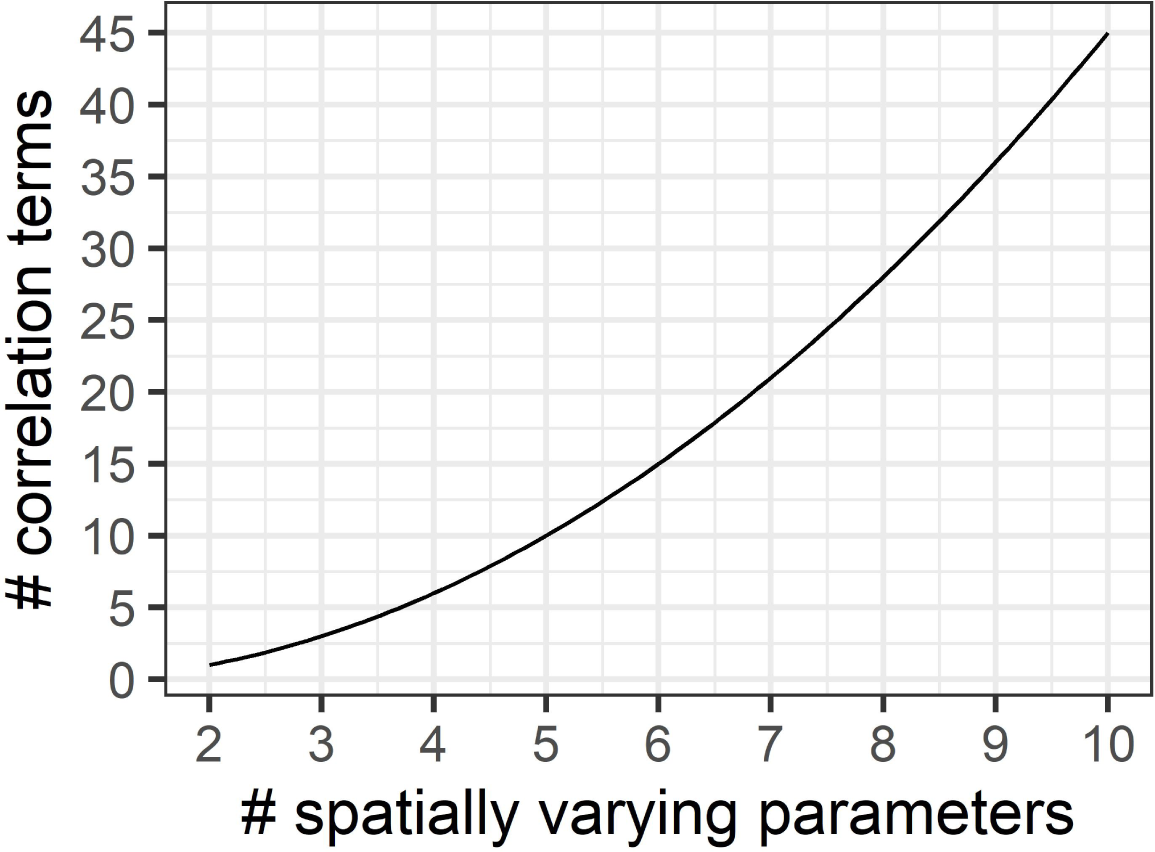
Model complexity grows exponentially with number of spatially varying parameters. We argue to keep models as simple as possible for both analytical and computational tractability.

As is clear in figure 2, the scale transition for temperature is extremely convex. Integration of fluxes over ecosystems with significant heterogeneity in temperature invokes substantial deviation from a mean-field model. For instance, at a CV of 0.2, the scale correction factor is 1.10, but by a CV of 0.5 it is 1.66. Obviously, the significance of this depends on the scale and heterogeneity over which an accurate flux model is desired. For a smaller footprint eddy covariance tower over a uniform habitat type, soil (and near surface) temperatures probably do not vary by much more than 20%, however this should be examined empirically. Regardless, our general mathematical analysis quantifies and clarifies exactly how the scale of variation influences the degree of the scale transition correction.

Notably, the only difference between the scale transition correction for first order and for reverse MM kinetics, is that in the latter, there would be additional correlation terms to consider, e.g. the correlation between temperature and *V*_*max*_, temperature and *k*_*h*_, as well as temperature and *C* and *MB*. In short, even under the assumption of linear first-order dynamics, ecological heterogeneity can induce a potentially substantial scale correction.

### Lessons for scientific inference

We close our discussion by considering the implications of the scale transition for advancing the state of biogeochemical modeling. Critically, the representation of non-linear (microbial driven) kinetics is a crucial modeling choice with large implications for long-term SOC forecasts. Traditional first-order PBMs avoid explicit representation of these kinetics, but nonetheless have worked well in practice. This state of affairs persists because both non-linear and linear kinetics are capable of representing coarse-scaled biogeochemistry reasonably well, at least in certain respects. Since first order kinetics are known to be a crude approximation, the crucial question for practice is not whether they are “true”, but rather whether their use involves significant, systematic information loss compared to non-linear microbial formulations. Fortunately, the scale transition offers several potential pathways to discriminate between these alternative model formulations.

As noted throughout, the dimensionless term *lambda* plays a critical role in linking the non-linear (michaelis-menten) kinetics to the first order kinetics. As *λ* increases, the non-linear kinetics converge to first order. Thus, in seeking to infer where the non-linear kinetic models provide substantial advantages, ensuring that *λ* is not too large (»1) is the first priority. Previous work Sihi et al. (2016) has approached this question theoretically, from first principles. Here, we point out that demonstrating substantial deviation from mean-field model when fitting non-linear kinetics to data is both a necessary and sufficient condition for infering that *λ* is not too large. Thus we recommend that time series of flux data be fit to both a first order and a non-linear kinetic model, where crucial covariates including subtrate (SOC), microbial biomass, and possibly environmental parameters such as temperature, have been measured sufficiently well to quantify the relevant variances and covariances. We recommend both predictive model evaluation (i.e. based on leave-one-out cross validation), in addition to metrics of model misfit or discrepancy (e.g. Sander Greenland’s S-value, Greenland (2019)).

In addition to the role of *λ*, our analysis also cleanly shows the contribution of other terms to the scale transition, and thus alternative metrics to assess. First and foremost, accounting for the spatial colocation of microbial biomass and substrate (in simplified form in equation 15) or alongside the various correlation terms between microbial biomass and kinetic/environmental factors (in full equation 9). In addition to fitting fully parameterized flux models, equation 9 suggests that low-hanging fruit may involve simply fitting statistical models including variations in microbial biomas, or colocation of microbial biomass and SOC, in explaining across-site variations in ecosystem respiratory fluxes (F). **A substantial role for either correlation of MB and C, or their variability, would constitute ipso facto evidence of the preferability of well-formulated non-linear kinetic models**. On the other hand, small roles for colocation, or evidence of large values of *λ* in practice would suggest minimal advantage to abandoning first order models in favor of more complex microbial models.

## Conclusions

Here, we have illustrated how the spatial scale transition can be expressed in dimensionless form, yielding insight into the systematic operation of Jensen’s Inequality in upscaling microbial decomposition kinetics. Our analysis has identified the central role of the dimensionless quantity *λ* - representing the ratio of mean-field microbial biomass over its half-saturation value - in governing the extent of the scale transition correction, expressed here in multiplicative form best facilitating comparison among systems. For somewhat simplified scenarios - such as restricting to spatial colocation of substrate and microbes - as *λ* → ∞, the mean-field correction goes to 0 and the model converges to first order.

This dual sense of convergence also provides opportunity to empirically test for the presence of significant non-linear microbial dynamics in upscaled field data: to the extent that upscaled fluxes deviate from the flux estimated at mean-field conditions, we have *ipso facto* evidence for the importance of formulating our biogeochemical models with these non-linear terms. Conversely, where there is close agreement between mean-field and upscaled fluxes, there are arguably stronger reasons for retaining first-order process model formulations.

In closing, we would like to point out how this mathematical analysis illustrates the challenge of scaling quite nicely. In the context of non-linear models, for each parameter that is allowed to vary in space, there is not only a new variance parameter, but a number of new covariance terms are induced, growing as the factorial of the number of varying parameters 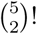 Thus, in the case of the 5 parameter function considered here, the full approximation has 5 mean field terms, 5 coefficients of variation, 10 correlation coefficients, and the dimensionless quantity *λ*.

Even with a maximally generic and simplified expression, fitting such models to field data still represents quite a challenge, especially while adequately accounting for and propagating uncertainty. Modelers and theoreticians should appreciate the complexity of the task at hand. Fortunately, our analysis has clearly identified a potentially robust route to managing model complexity: screen systematically for the importance of various correlations in explaining variations in fluxes. Accordingly, we recommend that research focus first upon spatial colocation of MB and C, which is readily measured, and then to thoughtfully and carefully expand models with additional terms as needed.

## Notes

### Competing Interest Statement

The authors have declared no competing interest.

